# PyVOL: a PyMOL plugin for visualization, comparison, and volume calculation of drug-binding sites

**DOI:** 10.1101/816702

**Authors:** Ryan H.B. Smith, Arvin C. Dar, Avner Schlessinger

## Abstract

**Motivation:** Binding pocket volumes are a simple yet important predictor of small molecule binding; however, generating visualizations of pocket topology and performing meaningful volume comparisons can be difficult with available tools. Current programs for accurate volume determination rely on extensive user input to define bulk solvent boundaries and to partition cavities into subpockets, increasing inter-user variability in measurements as well as time demands.

**Results:** We developed PyVOL, a python package with a PyMOL interface and GUI, to visualize, to characterize, and to compare binding pockets. PyVOL’s pocket identification algorithm is designed to maximize reproducibility through minimization of user-provided parameters, avoidance of grid-based methods, and automated subpocket identification. This approach permits efficient, scalable volume calculations.

**Availability:** PyVOL is released under the MIT License. Source code and documentation are available through github (https://github.com/schlessingerlab/pyvol/) with distribution through PyPI (bio-pyvol).

## 1 Introduction

Describing and predicting interactions between proteins and small molecule ligands, including natural substrates such as ATP and synthetic drugs, has been a major focus of computational structural biology. Many methods have been developed to identify putative binding sites, calculate physicochemical properties, and relate these properties to compound interactions (Kozakov, et al., 2015; Taylor, et al., 2002). These approaches are possible because of the high geometric, electrostatic, and hydrophobic complementarity between protein binding pockets and small molecules necessary for efficient, high-affinity interactions. While gestalt pocket properties are useful in explaining binding site interactions, druggable pockets often are composed of distinct subpockets with very different physicochemical properties. Decomposition of pockets into regions that correspond to likely small molecule pharmacophore-binding regions offers an intuitive improvement in predictive potential over exclusive use of average properties (Kuhn, et al., 2006).

Binding site prediction programs utilizing sequence and structure information can be based on a variety of approaches, including structural similarity to proteins with known binding sites, geometrical and energy calculations, or various combinations of these approaches (Zhang, et al., 2011). A subset of these methods such as POVME solely focus on volume calculation (Wagner, et al., 2017). A critical step in these methods that are designed for surface-exposed pockets is in distinguishing the pocket-associated volume from the surrounding bulk solvent, a highly non-trivial task that is normally performed either by the user, hindering reproducibility, or by overly conservative surface identification algorithms. Thus, more rigorous approaches for establishing this boundary are taken by programs designed to report meaningful volume calculations.

PyVOL is a unique pocket volume algorithm in several key ways. First, it only calculates solvent-accessible surfaces and ignores unrealistically small nooks or crannies in the protein’s Van der Waals surface. Second, PyVOL avoids using an underlying grid to perform pocket identification and volume determination. This eliminates the dependence of volume calculations on purely computational heuristics such as grid sampling, orientation, and origin. Third, PyVOL is designed to minimize the number of user-provided parameters in order to maximize inter-user replicability. While sufficient parameters are provided to permit user customization, parameters have been chosen as much as possible to have direct, obvious impacts on the results. Finally, PyVOL can perform automatic subpocket prediction through geometric partitioning on the basis of local concavity. Meaningful volume difference between pockets can be concentrated in one subpocket. Since volume calculations can be inherently noisy due to insufficiently sampled protein flexibility, minimizing the region over which differences will be calculated is essential to obtaining meaningful results. PyVOL’s automated functionality assists with identifying higher variance subpockets and in comparing them across large numbers of structures where manual specification becomes impractical.

## 2 Features

### 2.1 Binding Pocket Identification and Volume Calculation

PyVOL defines binding pockets as the volumes accessible to small probes and inaccessible to large probes (Oliveira, et al., 2014; Yu, et al., 2009). It distinguishes these binding pockets by first identifying the exterior surface of the protein that is exposed to bulk solvent. PyVOL determines a set of large-radius spheres tangent to the surface of the protein that completely encapsulate the protein. PyVOL calculates tangent spheres by clustering output from the program MSMS using DBSCAN (Ester, et al., 1996; Sanner, et al., 1996). All cavities within the bounded protein model that are accessible to a small-radius probes are then identified as potential pockets represented by collections of tangent spheres.

Pocket identification can be run in three modes: 1) it can identify all pockets, both surface-exposed and completely buried, that are larger than a user-specified volume, 2) it can return only the largest pocket, or 3) it can accept a user specification of a particular pocket of interest. Specification can be given as a coordinate, a surface residue, or a ligand. When a ligand is specified, the binding pocket can be augmented to include extensions of the ligand into the bulk solvent or limited to the region immediately surrounding the ligand. These options help maximize comparability between calculations of ligand and binding pocket size.

PyVOL stores pocket information both as the set of coordinates and radii of the identifying tangent spheres and as a three-dimensional triangulated mesh. Volumes are calculated using the Trimesh triangular mesh library. Unlike most volume calculation, this approach avoids explicit reliance on a grid for both pocket identification and volume calculation, eliminating variation in results caused by choice of grid sampling, orientation, and origin.

### 2.2 Subpocket Partitioning

Accurate subpocket estimation is useful for understanding substrate specificity in related proteins (e.g., glutamine vs glutamate transporters) as well as for designing chemicals targeting unique conformational states (e.g. inactive and active conformations in protein kinases) (Sonoshita, et al., 2018). Subpocket partitioning in PyVOL occurs through geometric recognition of locally concave regions within the overall pocket. Tangent spheres with radii varying between that of the small and large probes used in the first step are calculated within the binding pocket. Regions are identified by the largest internal tangent spheres while surface geometry is defined by the smallest internal tangent spheres. Hierarchical clustering propagates region assignments from larger spheres to smaller spheres. Spheres with a given radius are first clustered to the nearest large-radius sphere if sufficiently close. Unclustered, highly overlapping spheres with that same radius are then clustered with each other maximizing internal group overlap. Iteration of this process from the largest to the smallest spheres propagates region identification to the surface geometry.

A network capturing connectivity between regions is calculated through a metric correlated with volume overlap. Regions with insufficient total volume or insufficiently large maximum radii are merged into larger regions on the basis of the connectivity network. If the total number of regions exceeds an optional user-provided value, further consolidation occurs through the same process. Once each sphere is assigned to a region, overlap between regions is iteratively minimized. Final subpocket definition occurs through triangulation of the exterior hull for each region.

### 2.3 Pocket Comparison

Meaningful comparison between similar pockets or subpockets requires identification of analogous pockets, similar subpocket partitioning when relevant, and comparable bulk-solvent interface definitions. PyVOL allows independent specification of pocket positions for each protein being compared. Consistent partitioning of homologous pockets can usually be enforced through setting a reasonable maximum number of subpockets because this emphasizes overall pocket shape at the expense of minor differences within subpockets. While local variations in concavity can lead to identification of more or fewer subpockets, connectivity-based merging improves subpocket boundary consistency.

## 3 Examples

### 3.1 Catalytic Cleft Conformations for BRAF

The kinase BRAF has several well-characterized binding pocket conformations that are associated with its catalytic activity. These conformations are defined by the Asp-Gly-Phe (DFG)-motif and the αC-helix that can adopt “in” or “out” conformations (Fig 1). PyVOL captures the different subpockets available in three representative structures as shown in Fig. 1. Notably, the pocket partitioning identifies the new subpockets exposed by the conformational changes in BRAF.

**Figure 1.**
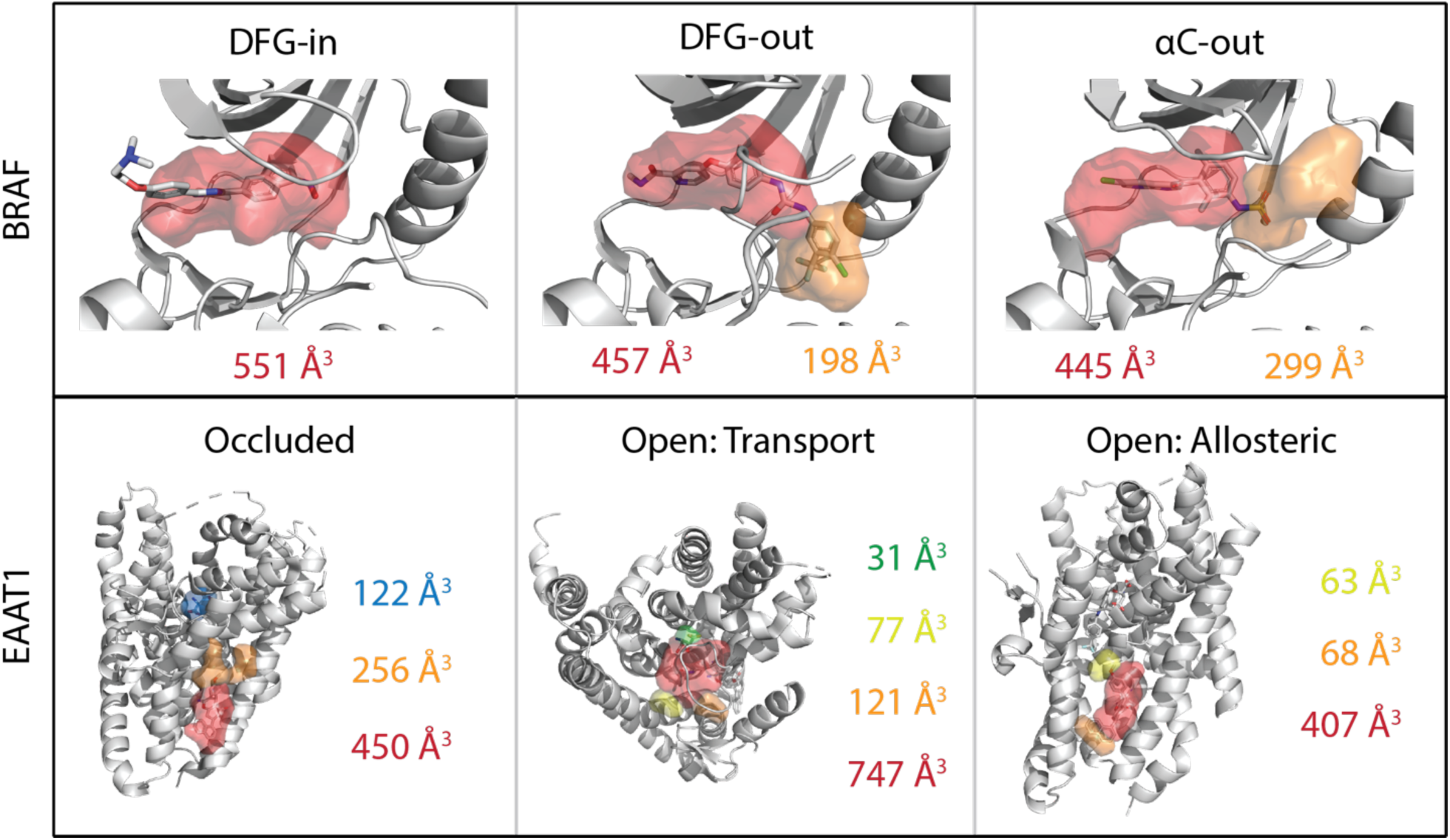
Examples of PyVOL’s output for three conformations of the kinase BRAF and three pockets found in two conformations of the glutamate transporter EAAT1. Calculated volumes for subpockets are shown for BRAF in the DFG-in (2BDF A), DFG-out (1UWH B), and αC-out (3C4C A) conformations. The pocket calculation for the DFG-in structure clearly shows the extension of the bound small molecule into solvent. EAAT1 is shown in the occluded (5LLM) and open (5MJU A) conformations with both transport (blue) and allosteric (all other colors) pockets shown.

### 3.2 Comparison of Open and Occluded Conformation Pockets of EAAT1

Three different pocket conformations have been observed for the Excitatory Amino Acid Transporter 1 (EAAT1) that capture different steps in the transport process. PyVOL can identify the substrate binding site including conformation-specific subpockets (Fig. 1).

## 4 Conclusion

PyVOL is a freely accessible PyMOL plugin to visualize, partition, and characterize binding pocket volumes. This package also includes programmatic and command-line interfaces to facilitate integration into bio-informatic pipelines. PyVOL requires Python 2.7-3.7, MSMS, Trimesh 2.36.29+, scipy 1.2.1, and scikit-learn 0.20.2 as well as their dependencies. The PyMOL plugin has been tested against PyMOL 2.0.0+. Source code, tutorials, and examples can be found on the github page.

